# Structural insights into human excitatory amino acid transporter EAAT2

**DOI:** 10.1101/2021.11.03.465409

**Authors:** Takafumi Kato, Tsukasa Kusakizako, Chunhuan Jin, LiLi Quan, Ryuichi Ohgaki, Suguru Okuda, Kan Kobayashi, Keitaro Yamashita, Tomohiro Nishizawa, Yoshikatsu Kanai, Osamu Nureki

**Affiliations:** Department of Biological Science, Graduate School of Science, The University of Tokyo, Tokyo, Japan; Department of Bio-system Pharmacology, Graduate School of Medicine, Osaka University, Osaka, Japan; Integrated Frontier Research for Medical Science Division, Institute for Open and Transdisciplinary Research Initiative (OTRI), Osaka University, Osaka, Japan

## Abstract

Glutamate is a pivotal excitatory neurotransmitter in mammalian brains, but excessive glutamate causes numerous neural disorders. Almost all extracellular glutamate is retrieved by the glial transporter, Excitatory Amino Acid Transporter 2 (EAAT2), belonging to the SLC1A family. However, in some cancers, EAAT2 expression is enhanced and causes resistance to therapies by metabolic disturbance. Despite its crucial roles, the detailed structural information about EAAT2 has not been available. Here, we report cryo-EM structures of human EAAT2 in substrate-free and selective inhibitor WAY213613-bound states. EAAT2 forms a trimer, with each protomer consisting of transport and scaffold domains. Along with a glutamate-binding site, the transport domain possesses a cavity, that could be disrupted during the transport cycle. WAY213613 occupies both the glutamate-binding site and cavity of EAAT2 to interfere with its alternating access, where the sensitivity is defined by the inner environment of the cavity. This is the first characterization of molecular features of EAAT2 and the selective inhibition mechanism, underlying structure-based drug design for EAAT2.

## Introduction

Amino acids are essential biomolecules for protein biosynthesis, metabolism and signal transduction, and the control of cellular amino -acid concentrations is quite important. Therefore, human cells have eleven discrete SoLute Carrier (SLC) families that transport various kinds of amino acids across the membrane^1,2^. The SLC1A family functions as a sodium-dependent symporter for the uptake of extracellular amino acids (Supplemental Fig. 1), and its members are categorized as Excitatory Amino Acid Transporters (EAATs; EAAT1–5 function as aspartate and glutamate transporters) and Alanine Serine Cysteine Transporters (ASCTs; ASCT1 and ACST2 function as neutral amino-acid transporters)^3^.

Higher functions of the mammalian central nervous system (CNS) are linked to complex neural activities, such as learning and memory^4^. In the CNS, glutamate, a principal excitatory neurotransmitter, stimulates ionotropic receptors to elicit the postsynaptic action potential via various ion fluxes, including calcium ions^5,6^, although excessive glutamate at synaptic clefts is associated with greater calcium influx. The intracellular accumulation of calcium ions is related to mitochondrial dysfunction and oxidative stress and induces neuronal cell death, known as excitotoxicity^7^. To protect neuronal cells from excitotoxity, the released glutamate is rapidly retrieved by transporters localized around the synaptic cleft. Especially, EAAT2 (also known as SLC1A2 or GLT-1) is highly expressed at the plasma membrane of glial cells and removes almost all (more than 90%) extracellular glutamate^8–10^. Therefore, EAAT2 plays a crucial role in the extracellular glutamate homeostasis. In accordance with its essential role, a deficiency in the EAAT2 transport activity is associated with serious diseases, including psychiatric and neurological disorders^11–17^.

Structural research on the SLC1A family has revealed the architectures and transport mechanisms of the archaeal homologues (Glt_ph_ and Glt_tk_)^18–25^ and four eukaryotic transporters (thermostabilized EAAT1, EAAT3, ASCT1 and ASCT2)^26–31^. SLC1A transporters are assembled into a trimer, with each protomer consisting of scaffold and transport domains to adopt a unique alternating access model, termed the “elevator-type mechanism”^20,32^. This model is operated by the rigid elevator-like movement of the transport domain to translocate substrates across the membrane. However, despite its pivotal role in the CNS, structural information about EAAT2 has not been reported. This information is particularly needed for pharmacological studies. Recent reports demonstrated that spider venom and a novel chemical compound function as “direct activators” to increase the transport activity of EAAT2 and provide neuroprotection against excitotoxicity^33–35^. In addition to neurological diseases, some kinds of tumors are related to the enhance expression of EAAT2, which is associated with resistance to a chemotherapeutic drug and endocrine therapies^36–38^. Since these resistances are clinical problems for patients, selective inhibitors of EAAT2 might be effective drugs for cancer therapies. Furthermore, highly selective inhibitors that can discriminate EAAT2 from other EAAT transporters will be useful for basic research to elucidate the physiological importance of EAAT2. Therefore, the structures of EAAT2 will provide molecular insights to facilitate the structural-based drug design of both activators and inhibitors.

In this work, we performed cryo-EM single particle analyses to determine the structures of human EAAT2. Our structures, together with transport assays and comparisons with other EAAT structures, provide insights into the molecular features of EAAT2 and the inhibitory mechanism of the highly selective inhibitor WAY213613.

### Structural determination and overall structure

We expressed full-length wild-type human EAAT2 (HsEAAT2) with a C-terminally-fused GFP tag (Supplemental Fig. 2) in mammalian Human Embryonic Kidney cells 293 (HEK293), and the recombinant proteins were purified with a GFP antibody^39^. For structural determination, purified HsEAAT2 proteins in glycol diosgenin (GDN) micelles were vitrified on grids, and 3,351 micrographs were recorded by a K3 camera. With the C3 symmetry imposed, we finally obtained the three-dimensional reconstruction map at the global resolution of 3.6 Å, based on the Fourier Shell Correlation (FSC) = 0.143 criterion (Supplemental Fig. 3). The densities of all transmembrane (TM) helices and β-strands were clearly observed (Supplemental Fig. 4), whereas the N- and C-termini, Ala110–Ser113, Lys148–Val162 and Lys194–Val229 were not detectable, suggesting that these regions are flexible.

HsEAAT2 forms a homotrimer (Fig. 1**a, b**), with each protomer consisting of eight TMs (TM1–8) and two helical hairpins (HP1 and HP2), which can be divided into transport and scaffold domains (Fig. 1**c, d**). The scaffold domains are located near the central symmetry axis and forms the trimer interactions, while the transport domains are located at the periphery of the trimer (Fig. 1**b**).

**Figure 1.**
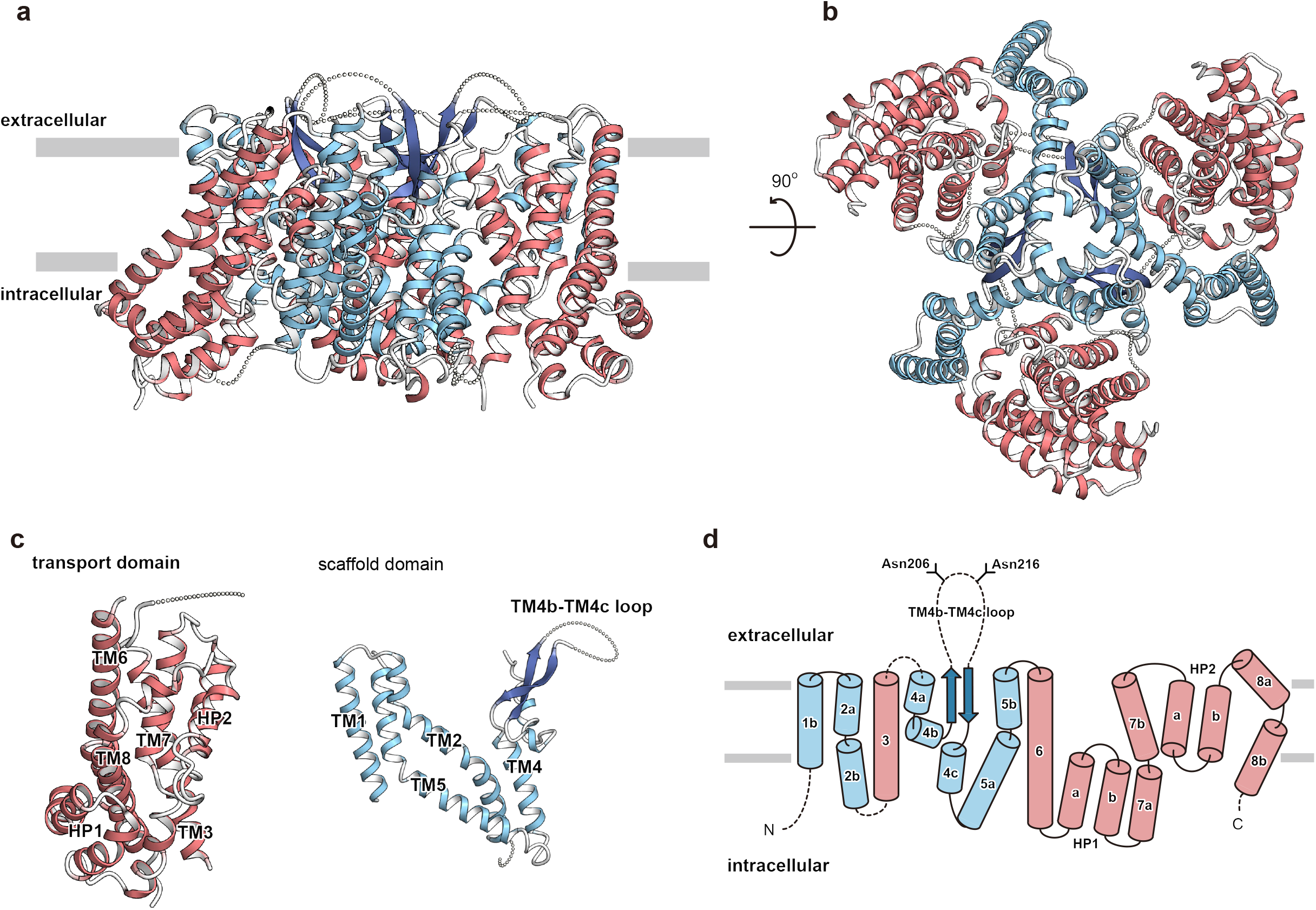
Overall structure of HsEAAT2. **a, b**, Overall structure of HsEAAT2, as viewed from (**a**) the membrane plane and (**b**) the intracellular side. **c**, The scaffold domain and the transport domain in one protomer are labeled. **d**, Topology diagram of HsEAAT2. The transport domain, the scaffold domain and the β-strands of TM4 are colored light red, light blue and dark blue, respectively. Two residues (Asn206 and Asn216) in the TM4b–c are putative glycosylation sites.

The transport domain consists of four TMs (TM3, TM6, TM7 and TM8), HP1 and HP2 (Fig. 1**c, d**), which comprise HP1a, b and HP2a, b, and the connecting HP1 and HP2 loops, respectively. Two hairpins are located on the domain interface, where HP2 contacts the scaffold domain in the membrane region, while HP1 is apart from the scaffold domain and located outside of the region (Fig. 2**a**). The transport domains partly protrude from the lipid bilayer by about 30 Å, and the putative glutamate-binding site is open toward the intracellular solvent (Fig. 2**a**). Therefore, the current structure represents the inward-facing state.

**Figure 2.**
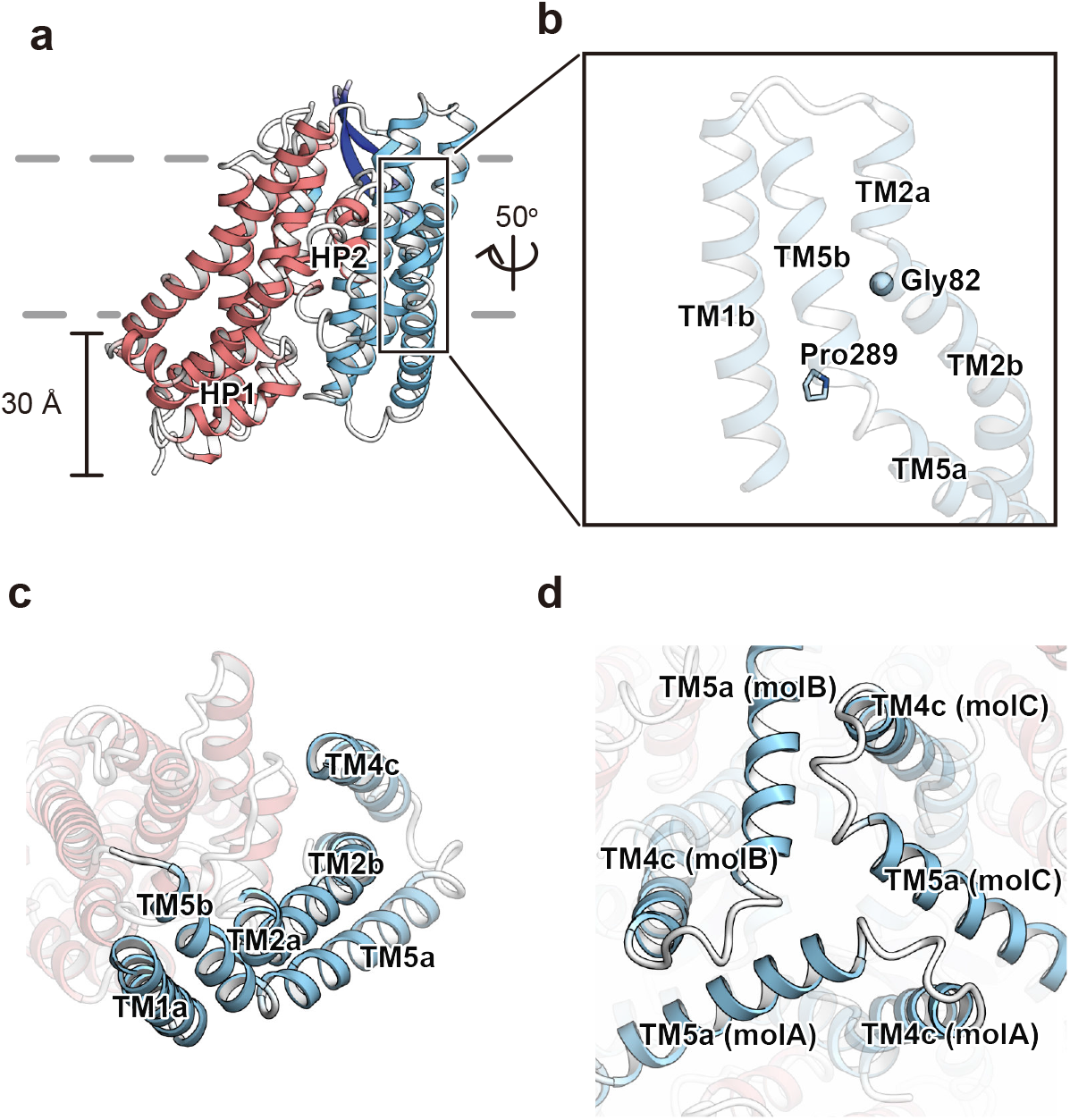
Protomer structure of HsEAAT2. **a**, Overall structure of the HsEAAT2 protomer, as viewed from the membrane plane. The distance indicates the protrusion of the transport domain from the lipid membrane. **b**, The “kink-induced” residues Gly82 (TM2) and Pro289 (TM5). Gly82 and Pro289 are located on TM2 and TM5, respectively. **c**, Interaction of TM1, TM2, TM4 and TM5 in the same protomer. **d**, The interactions among protomers on the intracellular side. “molA–molC” in brackets show each protomer.

The scaffold domain consists of four TMs (TM1, TM2, TM4 and TM5), and three of these TMs are divided into segments (TM2a, b, TM4a–c and TM5a, b) (Fig. 1**c, d**). TM2 and TM5 are kinked by Gly82 and Pro289 (Fig. 2**b**) and divided into two segments, and the extracellular segments of the two helices mediate the trimeric assembly (Fig. 2**b**-**d**). In agreement with our structural information, natural variants of Gly82 and Pro289 are associated with epileptic encephalopathies^40^, suggesting that these mutations hinder the molecular trimerization.

### Putative functions of TM4 loop and lipid-binding sites

All eukaryotic SLC1A transporters have a long loop between the TM4b and TM4c segments (TM4b–4c loop; Cys184–Asn241 in HsEAAT2), whereas such loops are not conserved in the archaeal homologues, Glt_ph_ and Glt_tk_ (Supplemental Fig. 2). In HsEAAT2, only the juxtamembrane portion of this loop forms antiparallel β-strands (Gln186–Lys193 and Lys231– Asp238) protruding from the scaffold domain, and the rest of the loop, including two potential glycosylation sites (Asn206 and Asn216), is completely disordered (Fig. 1**a, c, d**). The N206S mutation and a reduced glycosylation phenotype have been detected in neurological disorders^41,42^. Consistently, the N206S mutation hampers localization in the plasma membrane, hence causing a marked reduction in the EAAT2-mediated glutamate uptake^43^. Therefore, the β hairpin may structurally support the association of the flexible glycosylated loop with luminal and/or extracellular proteins during proper anterograde transports and the endocytic event which involves recycling to the plasma membrane, respectively.

The localizations and activities of transporters are affected by specific lipid environments, such as lipid composition and membrane thickness^44^. Some structural studies of SLC1A transporters reported lipid-binding sites^26–28^, and similarly, we observed two flat-shaped densities within each protomer. The density shapes and sizes suggested that they are likely derived from GDN and endogenous cholesterol (Supplemental Fig. 5). GDN is located between the transport and scaffold domains, and the cholesterol is on the cytoplasmic end of the scaffold domain, where it forms a π-π stacking interaction with Trp286 (TM5) (Supplemental Fig. 5**b, c**). EAAT2 tends to localize at cholesterol-rich microdomains, where cholesterol molecules are essential to sustain the transport activity^45,46^. Consistently, the reduction of EAAT2 activity in cholesterol-depleted membranes was reportedly observed in people with Alzheimer’s disease^47^. Since the GDN has a cholesterol-like moiety, we hypothesize that two native cholesterol molecules could be harbored in the GDN- and cholesterol-binding sites identified in the current structure, and probably contribute to the localization and/or structural stability of EAAT2. In particular, a similar cholesterol-binding-site of ASCT2 was observed near the cholesterol-binding site of HsEAAT2 (ref. ^27, 28^). Trp272 on ASCT2 TM5 (the corresponding residue of EAAT2 is Met283) also forms π-π stacking interactions with the cholesterol analogue. Trp286 is highly conserved among eukaryotic SLC1A transporters (Supplemental Fig. 2), and the cholesterol-binding sites of EAAT2 are located near the binding site of ASCT2, suggesting that the intracellular side of TM5 commonly participates in the cholesterol binding.

### Substrate-free state of the transport domain

In the transport domains of SLC1A transporters, amino acid-binding sites, which recognize aspartate, glutamate and neutral amino acids, are localized in between the HP1 and HP2 loops. Recent structural studies on the inward-facing and outward-facing states of ASCT2 proposed that only the HP2 loop functions as a gate for the binding sites, and this mechanism is termed the “one-gate elevator transport mechanism” ^28,29^. After the binding site closure by the HP2 loop, an elevator-like movement allows the transport of substrates across the membrane. While six highly conserved residues (Ser364, Thr401, Asp475, Arg478, Thr479 and Asn482 in HsEAAT2) constitute the putative glutamate-binding site in HsEAAT2, and the HP2 loop adopts an open conformation to allow access from the intracellular solvent, no density was observed at this site (Fig. 3**a**). Therefore, the current structure is likely to represent the substrate-free inward-open state. To evaluate the roles of those conserved residues, we measured the glutamate uptake activities of their point mutants, using *Xenopus* oocytes, which clearly showed that the transport activities of all mutants were essentially abolished (Fig. 3**b** and Supplemental Fig. 6**a**). These results indicate their indispensable roles in glutamate transport.

**Figure 3.**
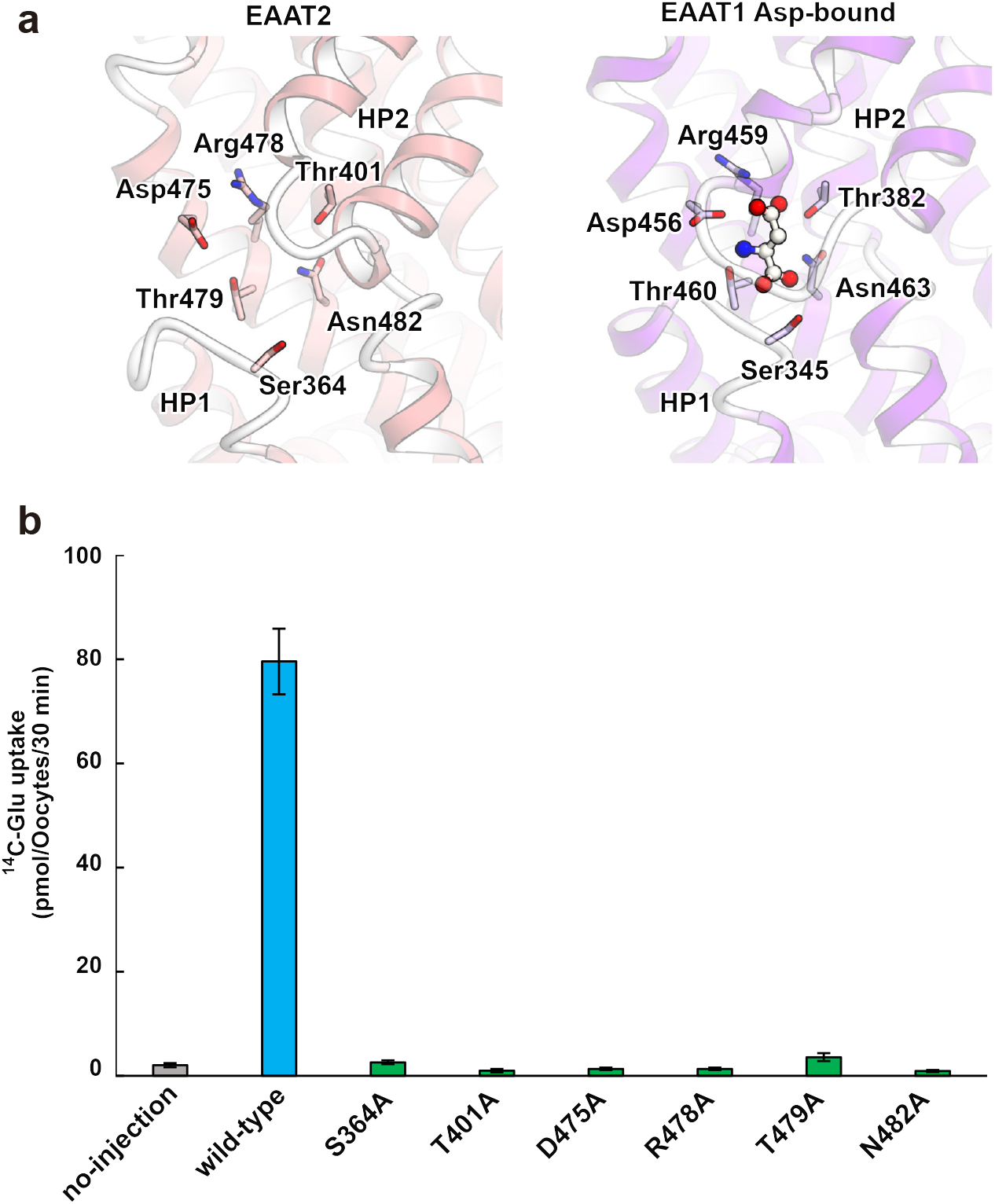
Glutamate-binding site. **a**, Comparison of the substrate-binding site in HsEAAT2 with the structure of the aspartate bound-state of EAAT1 (PDB ID 5LLU). **b**, Glutamate-uptake assay for point mutants, using *Xenopus* oocyte. Values are mean ± s.e.m. n= 6–10 technical replicates.

Amino acid transport by the SLC1A family members (EAATs, ASCTs and the archaeal homologues) is coupled with three sodium ions (Supplemental Fig. 1), and their binding sites have been clearly identified in previous structural studies^19,20,23,26,30^. In addition to sodium ions, EAATs utilize an extracellular H^+^ gradient, and its coupling mechanism was clarified in a recent report on EAAT3^30^. Firstly, in the occluded state, the HP2 loop functions as the gate to allow the binding of the transported amino -acid to the site. Next, the HP2 loop adopts the open conformation to release the amino -acid substrate, termed “IFS-Na^+^” in the previous work. In this state, the coupling of H^+^ neutralizes the charged glutamate residue (Glu405 and Glu374 in HsEAAT2 and EAAT3, respectively), whose protonation prevents the formation of a salt bridge with an arginine residue (Arg478 and Arg447 in HsEAAT2 and EAAT3, respectively) involved in the amino-acid substrate recognition. Upon the H^+^ release, the arginine residue adopts a different conformation to form the salt bridge with the deprotonated glutamate residue.

In our HsEAAT2 structure, the local structures of the three residues (Met398, Glu405 and Arg478) are similar to those in IFS-Na^+^ of EAAT3 (Supplemental Fig. 7). The p*K*_*a*_ value of Glu405 calculated by PROPKA program^48^ is 7.0, which is the same pH value in our purification. Considering the structural information and the p*K*_*a*_ value, Glu405 is probably in a transition between deprotonated and protonated forms, and does not stably form the salt bridge with Arg478. Therefore, our HsEAAT2 structure resembles the IFS-Na^+^ state of EAAT3. Since the transport domain adopts almost the same conformations in both the inward- and outward-facing states, behaving as a rigid body during the transport cycle^20,32^, a similar arrangement of the substrate-binding site could be observed in the outward-facing state of HsEAAT2.

### Inward-facing WAY213613-bound (IFS-WAY213613) state

The transport activities of EAATs are blocked by various inhibitors. For instance, TFB-TBOA is one of the strongest inhibitors for EAATs; it significantly suppresses the activities of not only EAAT2 but also other EAATs (IC_50_ values are 17, 22 and 300 nM for EAAT2, EAAT1 and EAAT3, respectively)^49^. Recently, WAY213613 was developed as a highly selective inhibitor of EAAT2 (IC_50_ values are 85, 5004 and 3787 nM for EAAT2, EAAT1 and EAAT3, respectively, in the HEK cell line)^50^. Among the available inhibitors, WAY213613 is the most potent and selective for EAAT2. To clarify the underlying inhibitory mechanism, we determined the cryo-EM structure of HsEAAT2 complexed with WAY213613 (Supplemental Fig. 8). The root mean square deviation value with the substrate-free state is 0.399 Å, indicating that the structures are very similar. Consistently, the transport domains represent the inward-facing states bound with WAY213613 (IFS-WAY213613 state), and the *F*_*o*_ – *F*_*c*_ omit map calculated by Servalcat program^51^ shows that WAY213613 is located in between HP1 and HP2 (Fig. 4**a, b**). WAY213613 is composed of two moieties: L-asparagine (LA) and 4-(2-bromo-4,5-difluorophenoxy) phenyl (BDP) moieties (Fig. 4**c**). The LA moiety is recognized by four residues (Thr401, Asp475, Arg478 and Thr479) at the glutamate-binding site (Fig. 4**a, d**), which are completely conserved among EAAT1–3 (Supplemental Fig. 2) and important for the glutamate transport (Fig. 3**b**). On the other hand, the BDP moiety is accommodated in a cavity located near the glutamate-binding site (Fig. 4**a, e**), formed by the end of HP2b, TM7 and TM8.

**Figure 4.**
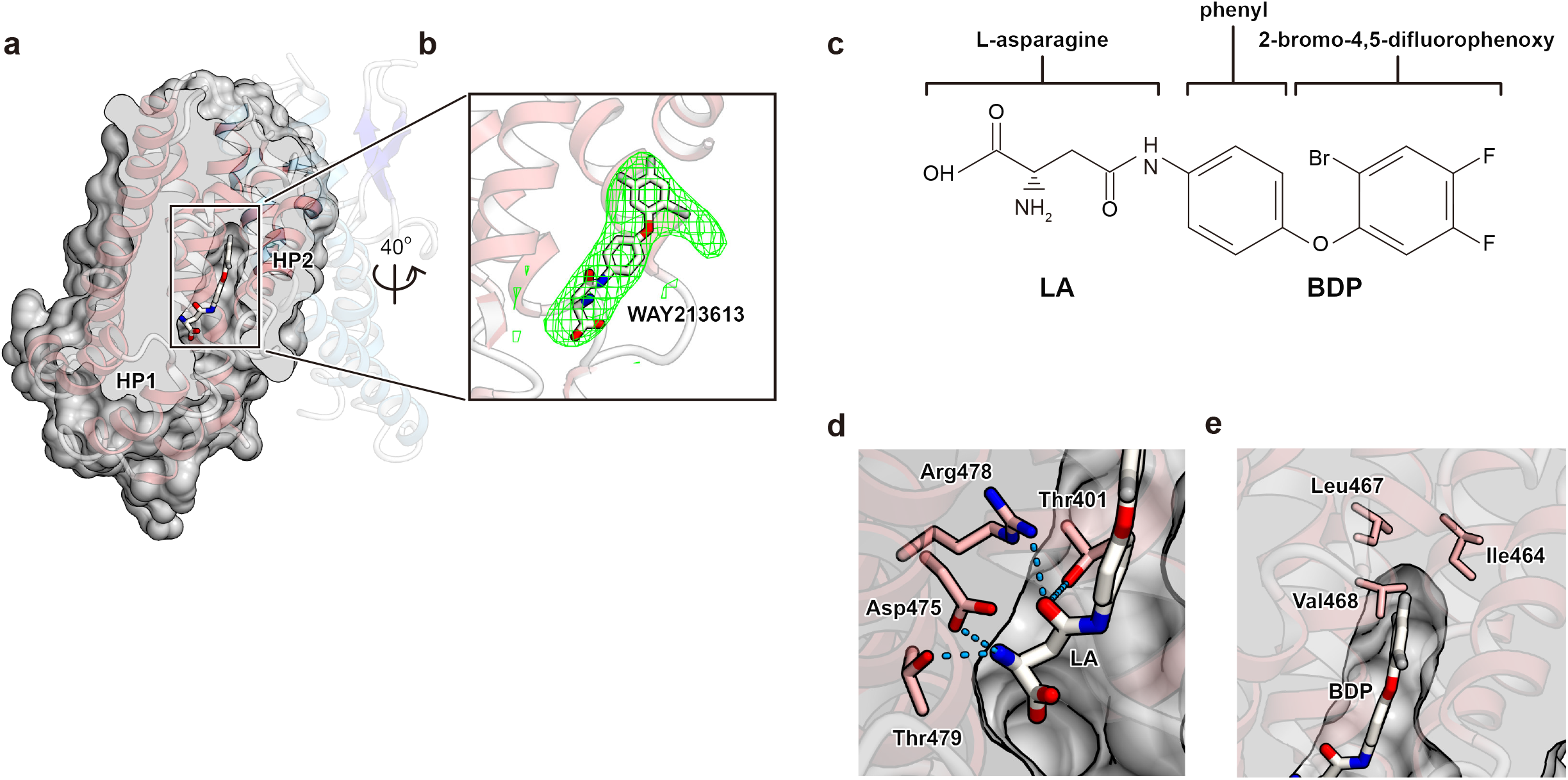
IFS-WAY213613 state. **a, b**, Structure of the IFS-WAY213613 state. **a**, Cut-away representation of the cavity with WAY213613. **b**, Close-up views of the WAY213613-bound site. The *F*_*o*_ – *F*_*c*_ omit map of WAY213613 is contoured at 4.0 s (normalized within mask), shown as a light green mesh. **c**, The structure of WAY213613. WAY213613 is composed of the L-asparagine (LA) moiety and the (2-Bromo-4,5-difluorophenoxy) phenyl (BDP) moiety. **d, e**, Recognition of the LA site of WAY213613. **d**, Substrate-binding site recognizing the LA moiety of WAY213613. **e**, Close-up view of the BDP moiety of WAY213613.

While the residues located around the BDP moiety are similarly conserved in EAATs (Supplemental Fig. 2 and 9), we found slight variations in three residues (Ile464, Leu467 and Val468), which are substituted with different sets of residues in EAAT1 and EAAT3 (Supplemental Fig. 2). As these residues are close to the tip of the 2-bromo-4,5-difluorophenoxy group of the BDP moiety (Fig. 4**c, e**), we supposed that the selectivity among EAAT1–3 may depend on the BDP moiety rather than the LA moiety. To verify our hypothesis, we designed point mutants, in which these residues are substituted with the corresponding residues of EAAT1 and/or EAAT3, and investigated the inhibitory effects of WAY213613 on the mutants, using *Xenopus* oocytes. All mutants showed similar transport activities to the wild type in the absence of WAY213613 (Fig. 5**a**), experimentally confirming that the mutations at the cavity have no or minimal effects on the glutamate uptake. The inhibition by WAY213613 was more or less affected by all three mutants (I464V, L467I and V468I). For instance, in the L467I mutant, although the IC_50_ value for EAAT2 is 130 nM in *Xenopus* oocytes^50^, the transport activity is hardly inhibited by WAY213613, showing almost the same level of glutamate uptake as the control even with the highest concentration (300 nM) of WAY213613 (Fig. 5**b** and Supplemental Fig. 6**b**). These results suggest that the cavity is closely related to the sensitivity of the EAAT subtypes to WAY213613.

**Figure 5.**
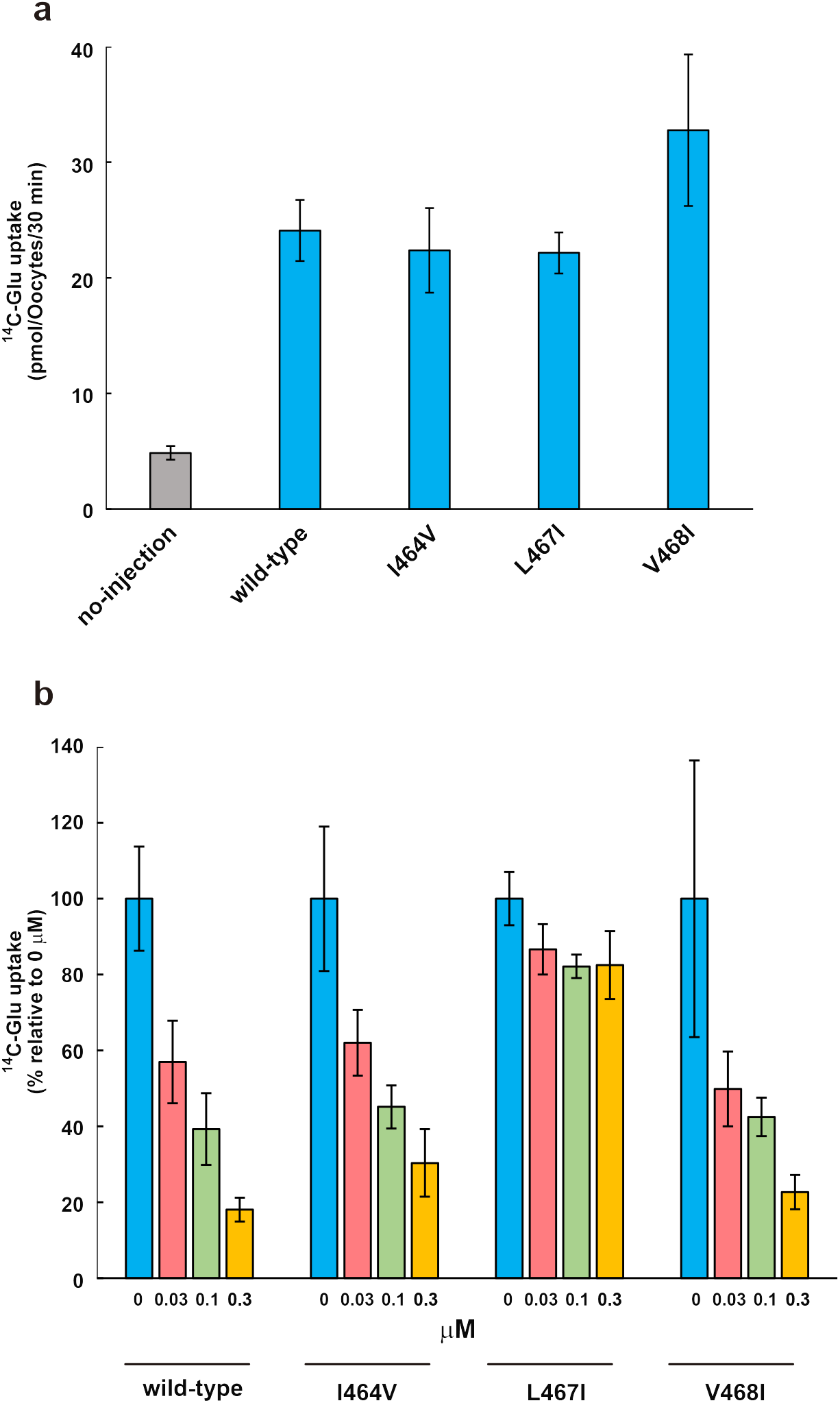
Sensitivity of WAY213613. **a**, Glutamate-uptake of each mutant, using *Xenopus* oocytes. **b**, Glutamate-uptake assay for point mutants treated with WAY213613. Values of vertical and horizontal axes are relative to each 0 μM and concentration points (0, 0.03, 0.1 and 0,3 μM) of WAY213613, respectively. Values are mean ± s.e.m. n= 6–10 technical replicates.

### Inhibitory mechanism of WAY213613

The LA moiety occupies the glutamate-binding site, where it probably competes with the glutamate binding, while the BDP moiety occupies the cavity near this binding site (Fig. 4**d, e**). A similar cavity is also present in IFS-Na^+^ of EAAT3 (Supplemental Fig. 10**a**). The mutation analysis demonstrated that the inhibitory effect of WAY213613 is largely diminished when Leu467 is substituted with the corresponding residues of EAAT1 and EAAT3 (isoleucine) (Fig. 5**a, b** and Supplemental Fig. 2). Considering the molecular superposition between the IFS-Na^+^ state of EAAT3 and the IFS-WAY213613 state of EAAT2, the γ^2^ carbon of isoleucine would narrow the cavity and sterically interfere with the binding of the 2-bromo-4,5-difluorophenoxy group of the BDP moiety (Fig. 6**a**). Altogether, the selectivity of WAY213613 is determined by the different local environments with in the cavities among EAATs. In EAAT1 and EAAT3, slight changes would prevent the proper accommodation of the BDP moiety and thus lead to the lower affinity for WAY213613.

The conformation of the HP2 loop in the IFS-WAY213613 state is very similar to that in the substrate-free state. The comparison among this IFS-WAY213613 state of EAAT2 and the Asp-bound states of EAAT1 and EAAT3 revealed that the end of HP2b slightly moves towards TM7 and TM8 upon aspartate binding, which accompanies the closing of the HP2 loop gate (Supplemental Fig. 10**b**). As the BDP moiety is likely to sterically interfere with the movements of HP2b and the HP2 loop, WAY213613 probably prevents the HP2 loop gating of EAAT2. Therefore, the LA and BDP moieties of WAY213613 play distinct roles in the EAAT2 inhibition: by competing with the glutamate binding and sterically preventing the HP2 loop gating, respectively. Closure of the HP2 loop is essential for the elevator-like movement of the transport domain^29^, suggesting that the BDP moiety locks the conformation of the HP2 loop to suspend the transport cycle. Whereas our structure complexed with WAY213613 represents the inward-facing state, the presence of WAY213613 in the extracellular solution clearly affected both the wild type and mutants in the oocyte assay (Fig. 5**b**). As the transport domains of both the inward- and outward-facing states adopt almost the same conformations and hence behave as rigid bodies during the transport cycle^20^, WAY213613 could bind the transport domain in the outward-facing state (Fig. 6**b**).

**Figure 6.**
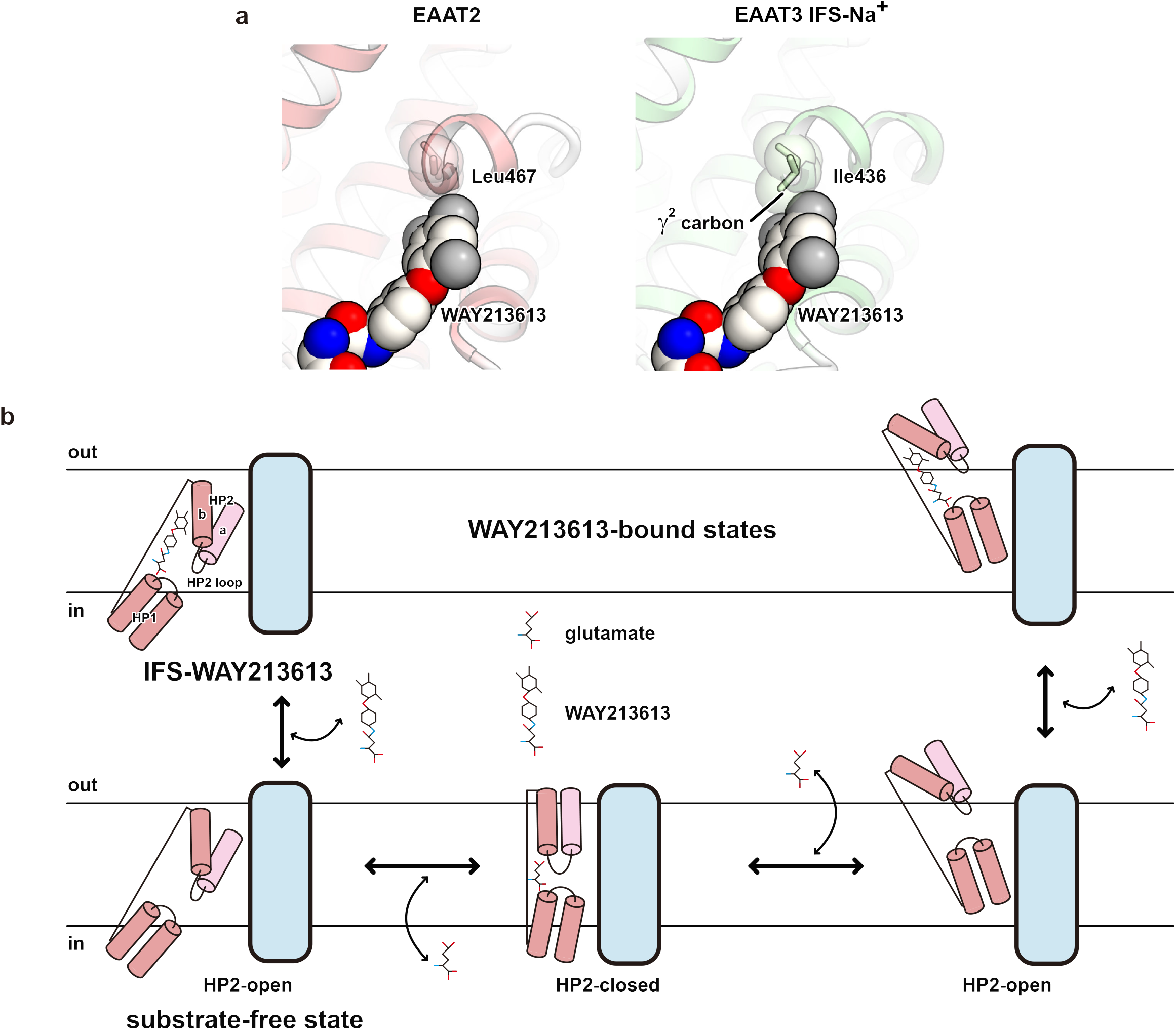
Inhibition mechanism by WAY213613. **a**, Close-up view of the IFS-WAY213613 state and molecular superposition of EAAT3 IFS-Na^+^. WAY213613, Leu467 (HsEAAT2) and Ile436 (EAAT3) are shown as CPK models. **b**, In the HP2-open configuration of the outward-facing state, extracellular glutamate is accessible at the substrate-binding site, and subsequently, HP2 is closed to transport glutamate into the intracellular solvent (bottom model). WAY213613 inhibits the movement of the HP2 loop (top model).

### Conclusion

Despite the essential role of glutamate as a neural transmitter in the CNS, excessive glutamate is toxic to neurons. EAAT2 clears almost all extracellular glutamate to maintain a low concentration, and dysfunction of the transporter leads to numerous neurological disorders. Therefore, extensive research has been conducted to understand the mechanism of EAAT2. In this work, we determined the structures of EAAT2 in the substrate-free and the inhibitor-bound states, and clarified the structural basis for its molecular features. Especially, the WAY213613-bound structure revealed the characteristic inhibitory mechanisms by its two moieties.

We observed the densities of cholesterol-related molecules in the present HsEAAT2 structure. A large portion of EAAT2 in the plasma membrane prefers to be localized at lipid rafts, reportedly affecting glutamate transport^45^. The detailed mechanism explaining why EAAT2 requires cholesterol molecules is still unclear. However, a recent report found that cholesterol molecules enhance the transport rate of ASCT2, showing that these molecules facilitate the elevator-like movement of the transport domains^52^. Since the conformational change of the transport domain induces the local membrane deformation^24^, the cholesterol may change the transport dynamics by altering the membrane properties^53^, to facilitate the uptake of extracellular glutamate. Therefore, our structural information will help to clarify the relationships between EAAT2 and lipid molecules that enhance the proper localization and activity of EAAT2. According to the inhibitory mechanism of WAY213613, more selective inhibitors for EAAT2 or other EAAT subtypes could be developed by modifying the moiety corresponding to BDP in WAY213613. Such selective inhibitors will illuminate more details of the physiological roles of EAAT2 in extracellular glutamate homeostasis, and may pave the way for future research and strategies for cancer therapies.

## Acknowledgements

We thank all of the members of the Nureki and Kanai laboratories, R. Danev and M. Kikkawa for setting up the cryo-EM infrastructure and K. Ogomori for technical assistance. This work was supported by a grant from JSPS KAKENHI grants 16H06294 (O.N.), and by the Platform Project for Supporting Drug Discovery and Life Science Research (Basis for Supporting Innovation Drug Discovery and Life Science Research (BINDS)) from the Japan Agency for Medical Research and Development (AMED) under grant no. JP19am0101115.

## Contributions

T. Kato purified and performed the cryo-EM trials of EAAT2, determined the structure, and planned the mutational analyses, under the supervision of T. N, Y. K. and O. N. K. K., T. Kusakizako, K. Y. and T. N. assisted with the EM image data collection, the data analyses and model building. R.O. designed mutants for transport measurements and expression analyses in *X. laevis*, and C. J., L. Q. and S. O. performed glutamate uptake analyses, localization analyses and western blotting, respectively, under the supervision of Y. K., T. Kato, T. Kusakizako, T. N. and O. N. wrote the manuscript, with feedback from all of the authors. Y. K. and O. N. supervised the research.

## Methods

### Purification of HsEAAT2

The sequence encoding the full-length human EAAT2 isoform 1 (SLC1A2; Uniprot ID P43004) was amplified from a human brain complementary DNA library (Zyagen) and inserted into the pEG BacMam vector^54^, with the C-terminally fused tobacco etch virus (TEV) protease cleavage site, enhanced green fluorescent protein (eGFP) and His_8_-tag. Baculoviruses were generated in *Spodoptera frugiperda* Sf9 cells, using the Bac-to-Bac system (Invitrogen).

HEK293 GnTI^-^ cells were grown and maintained in FreeStyle 293 medium (Gibco) supplemented with 2% fetal bovine serum, with 8% CO_2_ under humidified conditions. P2 baculoviruss were added to at a density of approximate 3.0 × 10^6^ cells/mL. Cells were cultured at 37°C for 24 hours. To boost overexpression, sodium butyrate was added at a final concentration 10 mM, and cells were cultured at 30°C for 48 hours. Cells were collected by centrifugation at 5,000 g for 6 minutes, resuspended in Buffer A (50 mM HEPES-NaOH, pH 7.0, 300 mM NaCl and 10% glycerol) and disrupted by probe sonication for 3 minutes. The debris was removed by centrifugation (8,000g, 10 min, 4°C), and subsequently membrane was collected by ultracentrifugation (125,000g, 1 h, 4°C). Membrane pellet was resuspended in Buffer A, homogenized in a glass homogenizer and store at −80°C.

All purification procedures were performed at 4°C. The membrane fraction was solubilized in Buffer A containing 1.0% lauryl maltose neopentyl glycol for 1 hour. After removing the insoluble material by ultracentrifugation (125,000g, 30 min, 4°C), the supernatant was incubated with CNBr-Activated Sepharose 4 Fast Flow beads (GE Healthcare) coupled with an anti-GFP nanobody^39^ resin for 3 hours. The resin was poured into an open column and washed with 15 column volumes of Buffer A containing 0.2% glycol diosgenin (GDN). The resin was mixed with both TEV protease and Buffer A containing 2.0 mM dithiothreitol (DTT) and 0.2% GDN overnight. The resin was poured into the open column, and elution was collected and concentrated for subsequent gel filtration chromatography (Superose 6 Increase 10/300 GL, GE Healthcare) in SEC Buffer (50 mM HEPES-NaOH, pH 7.0, 300 mM NaCl, 2.0 mM DTT and 0.05% GDN). The fraction containing the HsEAAT2 proteins was pooled and concentrated to 5-7 mg mL^-1^ using an Amicon Ultra Filter (MWCO 100 kDa). In the structural analysis of the IFS-WAY213613, the HsEAAT2 proteins was incubated with 1.0 mM WAY213613 (Tocris) for 1 hour on ice.

### Sample vitrification and cryo-EM data acquisition

The purified HsEAAT2 proteins was applied onto a freshly glow-discharged Quantifoil holey carbon grid (R1.2/1.3, Cu/Rh, 300 mesh), blotted for 4 seconds at 4 °C in 100% humidity and plunge-frozen in liquid ethane by using Vitrobot Mark IV (Thermo Fisher Scientific).

The grids were transferred to a Titan Krios G3i microscope (Thermo Fisher Scientific), running at 300 kV and equipped with a Gatan BioQuantum Energy Filter (GIF) and a Gatan K3 direct electron detector in the electron counting mode. Imaging was obtained at a nominal magnification of 105,000×, corresponding to a calibrated pixel size of 0.83 Å/pix. Each image was dose-fractionated to 63 (substrate-free state) or 48 (IFS-WAY213613) frames at a dose rate of 15 e^-^ per pixel per second, to accumulate a total dose of ∼50 e^-^ Å^-2^. The data were automatically acquired by the image shift method using the SerialEM software, with a defocus range of −0.8 to −1.6 μm.

### Data processing and model building

Image processing was performed in RELION-3.1 (ref. ^55^). The movie frames were aligned in 4 × 4 patches and dose-weighted with RELION’s implementation of the MotionCor2 algorithm^56^, and defocus parameters were estimated by CTFFIND 4.1 (ref. ^57^). In the apo-state, template-based auto-picking was performed with the 2D class averages of a few hundred manually picked particles as templates. A total of 1,090,865 particles were extracted in 3.1125 Å pix^-1^. These particles were subjected to one round of 2D classification. The initial 3D reference map was generated in RELION. Subsequently, 674,221 good particles were further classified in 3D classification in C3 symmetry. Finally, 212,554 particles were re-extracted in the pixel size of 1.55625 Å pix^-1^ and refined in C3 symmetry. The resulting 3D models and particle sets were subjected to per-particle defocus refinement, Bayesian polishing^58^, CTF refinement, and 3D refinement. The final 3D refinement and post-processing yielded map with global resolutions of 3.6 Å, with the gold standard Fourier shell correlation criteria (FSC = 0.143). In the WAY213613-bound state, a total of 831,890 particles were extracted in 3.1125 Å pix^-1^. Subsequently, similar processes were performed, and the overall gold-standard resolution was 3.49 Å. The initial structural model of HsEAAT2 was built by Phyre2 program^59^, based on human ASCT2 (PDB 6GCT). After model fitting by MOLREP program^60^, the models were manually readjusted using COOT^61^, and then refined using PHENIX^62^. The model and restrain information of WAY213613 were generated by eLBOW program^63^. Finally, the models and the *F*_*o*_ – *F*_*c*_ omit map of WAY213613 were refined and generated, respectively, by Refmac5 (ref. ^64^) using Servalcat^51^ under C3 symmetry constraints. The figures depicting the molecular structures were prepared using CueMol (http://www.cuemol.org/).

### Transport measurements and expression analyses in *X. laevis* oocytes

EAAT2 coding sequence was amplified by PCR from pEG BacMam-HsEAAT2 with the following primer pair: forward 5’-GGG<u>GGATCC</u>*GCCACC*ATGGCATCTACGGAAGGTGCCAAC-3’ (BamHI site underlined, Kozak sequence in italics) and reverse 5’-CCC<u>GAATTC</u>TCATTTCTCACGTTTCCAAGGTTCTTC-3’ (EcoRI site underlined). The PCR product was cloned into pcDNA3.1(+) (Invitrogen) at BamHI/EcoRI sites to obtain pcDNA3.1(+)-HsEAAT2. Mutations were introduced by whole-plasmid PCR using PrimeSTAR MAX DNA polymerase (Takara) according to the manufacturer’s protocol. Amino acid substitutions to alanine (for S364A, T401A, D475A, R478A, T479A and N482A), valine (for I464V) and isoleucine (for L467I and V468I) were performed by altering the corresponding codons into GCA, GTA and ATC, respectively.

Transport measurements and expression analyses in *X. laevis* oocytes were performed as described previously with minor alterations^65^. cRNAs of EAAT2 (wild-type and mutants) were synthesized *in vitro* from EcoRI-linearized plasmids, polyadenylated, and injected into defolliculated oocytes (25 ng/oocyte). The uptake measurements were performed 3d after injection, in ND96 buffer (96 mM NaCl, 2 mM KCl, 1.8 mM CaCl_2_, 1 mM MgCl_2_, 5 mM HEPES, pH 7.4) containing 50 μM of ^14^C-Glutamate (24.2 Ci mol-1, ARC, St. Louis, U.S.A.) for 30 min at room temperature. For inhibition experiments, the uptake of 10 μM of ^14^C-Glutamate was measured with or without the indicated concentrations of WAY213613 (Tocris, Bio-Techne, US). Expression of EAAT2 in total membranes was analyzed by immunoblotting. Anti-EAAT2 antibody (sc-365634, 1:400, Santa Cruz Biotechnology) and peroxidase goat anti-mouse IgG (AB_10015289, 1:10,000, Jackson ImmunoResearch) were used. Localization of EAAT2 was analyzed by immunofluorescence on paraffin sections. Antigen retrieval was performed with citrate buffer (0.01 M, pH6.0) at 121 °C for 5 min. Anti-EAAT2 antibody (sc-365634, 1:200, Santa Cruz Biotechnology) and Alexa Fluor 568-conjugated anti-mouse IgG (A11031, 1:2000, Invitrogen) were used. Images were acquired using a fluorescence microscope (BZ-9000, Keyence) equipped with a ×40 objective lens (CFI Plan Apo λ, numerical aperture 0.95, Nikon).

## Data availability

The cryo-EM density maps and atomic coordinates have been deposited in the Electron Microscopy Data Bank (EMDB). The accession codes for the maps are EMD-32098 (the substrate-free state) and EMD-32097 (the IFS-WAY213613 state). The PDB accession codes for the coordinates are 7VR8 (the substrate-free state) and 7VR7 (the IFS-WAY213613 state).

---

**Cryo-EM data collection, refinement and validation statistics**

|  | HsEAAT2 substrate-free state<br>(EMDB-32098)<br>(PDB 7VR8) | HsEAAT2 IFS-WAY213613<br>(EMDB-32097)<br>(PDB 7VR7) |
| --- | --- | --- |
| **Data collection and processing** |  |  |
| Magnification | × 105,000 | × 105,000 |
| Voltage (kV) | 300 | 300 |
| Electron exposure (e-/Å<sup>2</sup>) | 50 | 50 |
| Defocus range (μm) | -0.8 to -1.6 | -0.8 to -1.6 |
| Pixel size (Å) | 0.83 | 0.83 |
| Symmetry imposed | C3 | C3 |
| Initial particle images (no.) | 1,090,865 | 831,890 |
| Final particle images (no.) | 212,554 | 527,996 |
| Map resolution (Å) | 3.58 | 3.49 |
| FSC threshold | 0.143 | 0.143 |
| **Refinement** |  |  |
| Initial model used (PDB code) | 6GCT |  |
| Model composition in the asymmetric unit |  |  |
| Non-hydrogen atoms | 3,012 | 3,012 |
| Protein residues | 402 | 402 |
| Ligands<sup>a</sup> | 2 | 3 |
| Average *B* factors (Å<sup>2</sup>) |  |  |
| Protein | 137.6 | 129.3 |
| Ligand | 165.3 | 166.3 |
| R.m.s. deviations |  |  |
| Bond lengths (Å) | 0.016 | 0.015 |
| Bond angles (°) | 1.98 | 1.92 |
| Validation |  |  |
| MolProbity score | 2.21 | 1.17 |
| Clashscore | 13.31 | 3.17 |
| Poor rotamers (%) | 2.44 | 0.91 |
| Ramachandran plot |  |  |
| Favored (%) | 95.94 | 97.72 |
| Allowed (%) | 4.06 | 2.28 |
| Outlier (%) | 0 | 0 |

<sup>a</sup> Ligands include cholesterol, head moiety of GDN and WAY213613

