## Supplemental Figures for "Structural insights into human excitatory amino acid transporter EAAT2"

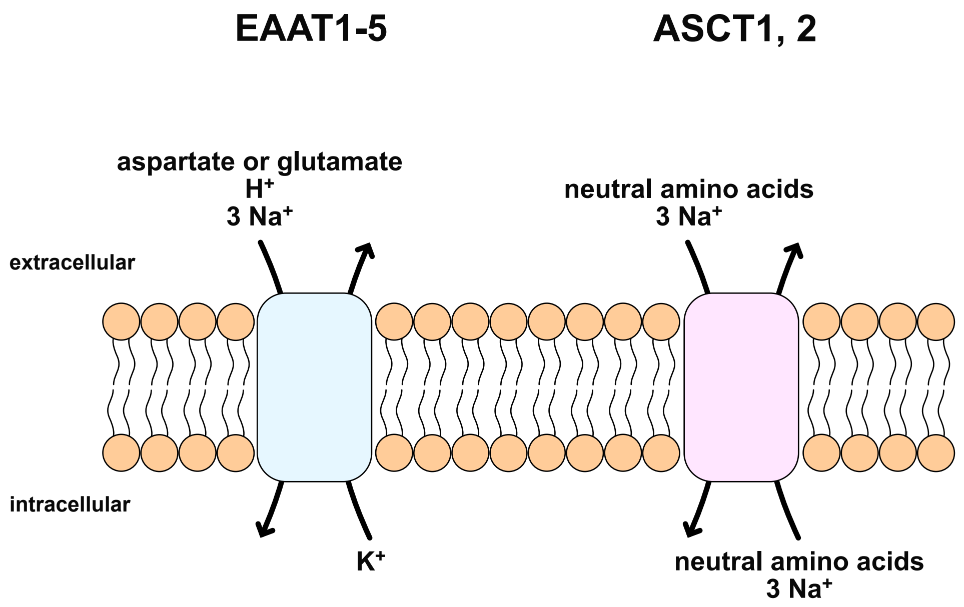


**Supplementary Figure 1 │ Substrates and coupling ions of SLC1A transporters**

Amino acid transport by all SLC1As is coupled to three extracellular sodium ions. In addition to sodium ions, extracellular proton and intracellular potassium ions are utilized for the transport cycle of EAATs.

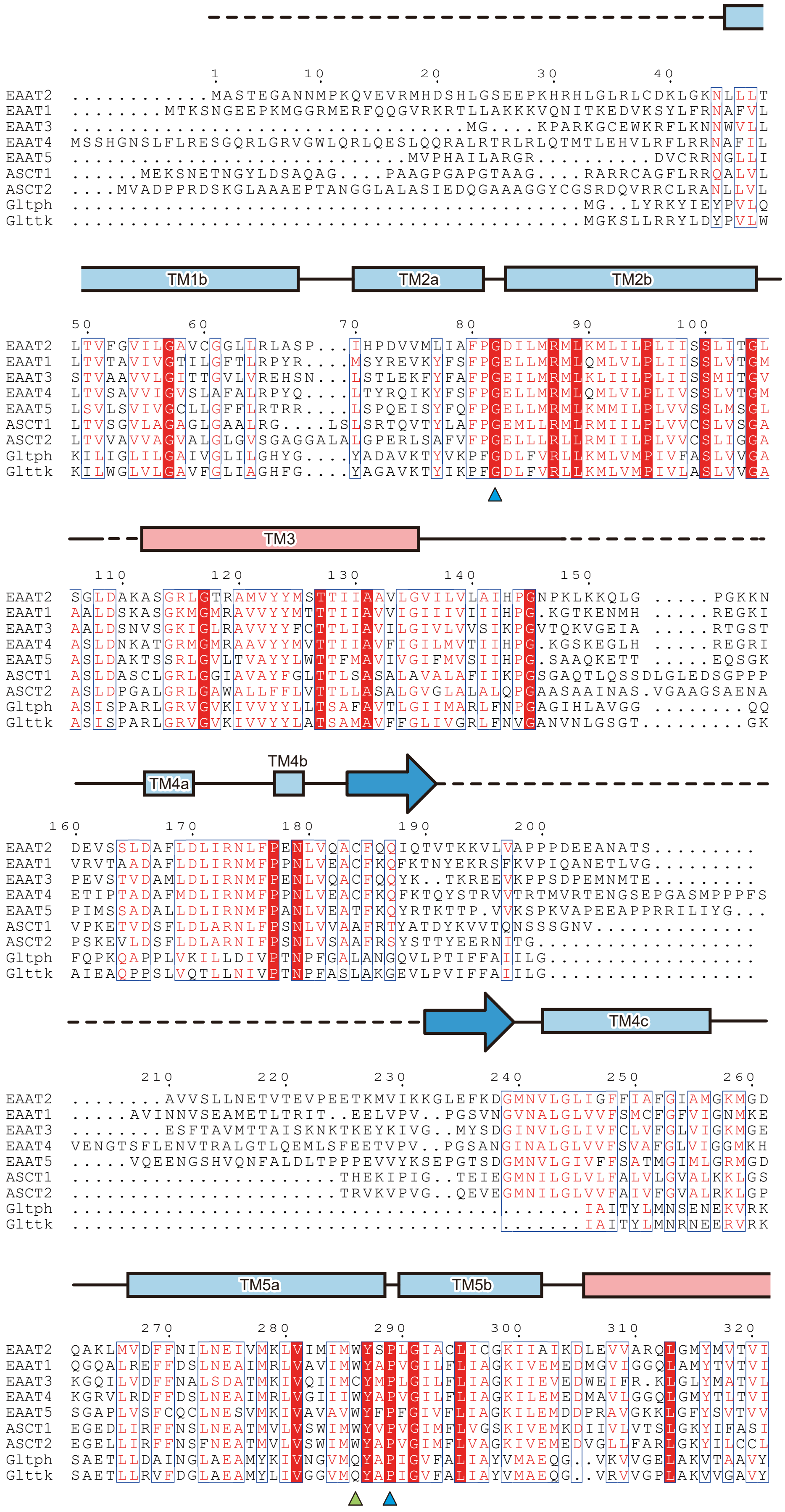

TM1b
TM2a
TM2b
TM3
TM4a
TM4b
TM4c
TM5a
TM5b
EAAT2
EAAT1
EAAT3
EAAT4
EAAT5
ASCT1
ASCT2
Gltph
Glttk

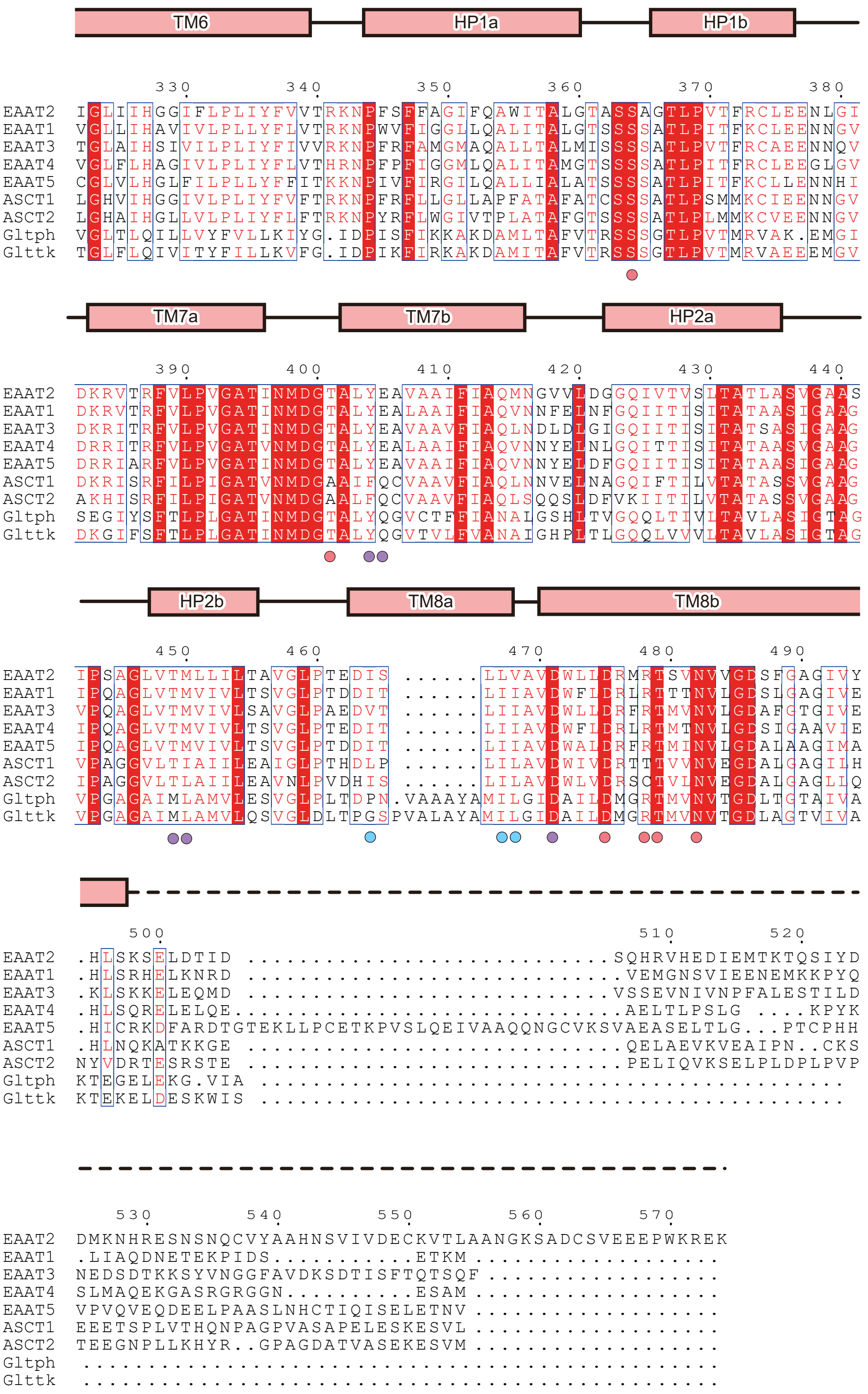


**Supplementary Figure 2 | Multiple amino acid sequence alignment of EAAT2**

Sequence alignment of *Homo sapiens* EAAT2 (HsEAAT2: UniProt ID P43004), *Homo sapiens* EAAT1 (P43003), *Homo sapiens* EAAT3 (P43005), *Homo sapiens* EAAT4 (P48664), *Homo sapiens* EAAT5 (O00341), *Homo sapiens* ASCT1 (P43007), *Homo sapiens* ASCT2 (Q15758), *Pyrococcus horikoshii* $Glt_{ph}$ (O59090) and *Thermococcus kodakarensis* $Glt_{tk}$ (Q5JID0). The secondary structure of HsEAAT2 is indicated above the sequence. The α-helices, β-strands and disordered regions are indicated by cylinders, arrows and dashed lines, and the scaffold domain and the transport domain are colored light blue and light red, respectively. At the scaffold domain, blue and green triangles indicate kink-inducing residues and Trp286, respectively. At the transport domain, red, purple and blue circles indicate residues of the glutamate-binding site, the cavity the tip around WAY213613, respectively.

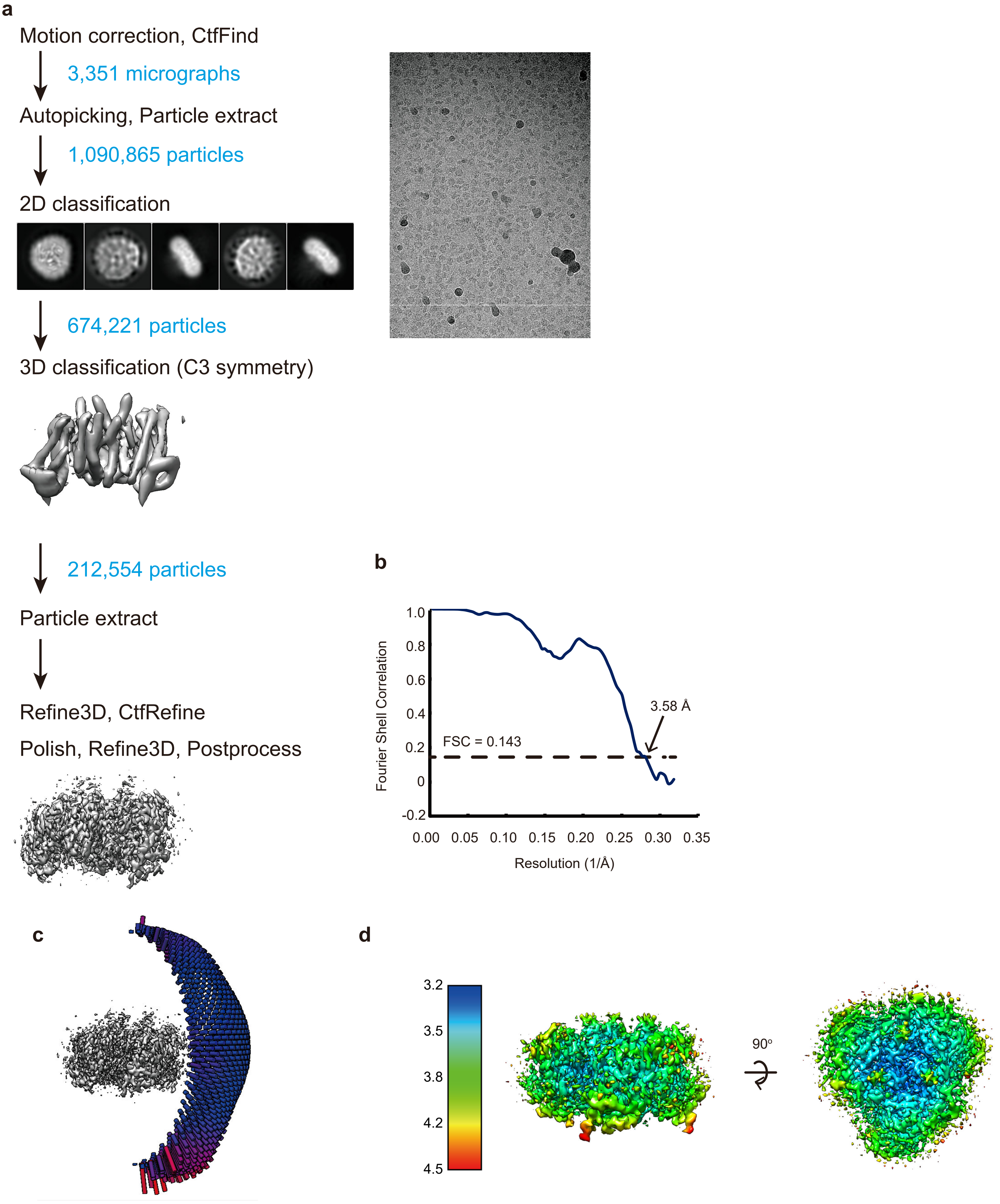


**Supplementary Figure 3│cryo-EM analysis of apo-state HsEAAT2**

**a,** Flow chart of cryo-EM data processing of HsEAAT2 apo-state structure. **b**, Fourier Shell Correlation (FSC) curve of the final 3D reconstruction model calculated using "relion_postprocess" with masked marked 3.98 Å resolution, corresponding to the FSC = 0.143 gold standard cut-off criterion. **c**, Angular distribution plot of particles included in the final 3D reconstruction of the HsEAAT2 apo-state structure, with C3 symmetry imposed. **d**, Local resolution of the HsEAAT2 apo-state structure, estimated by RELION.

**a**

Scaffold domain

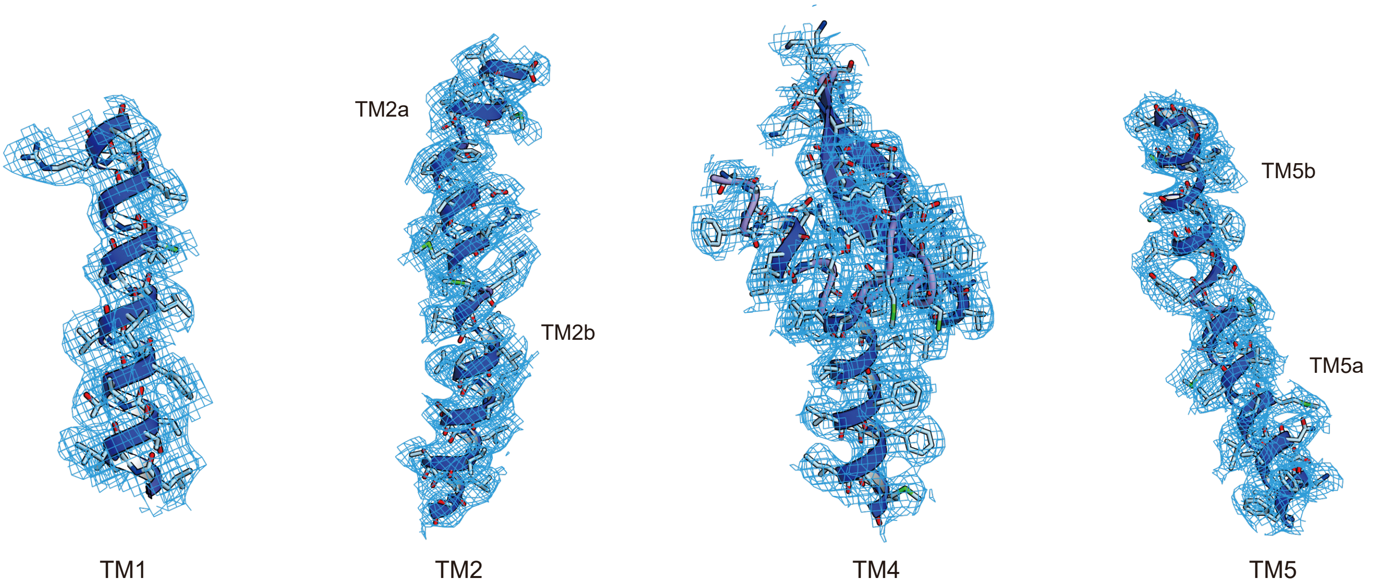


**b**

Transport domain

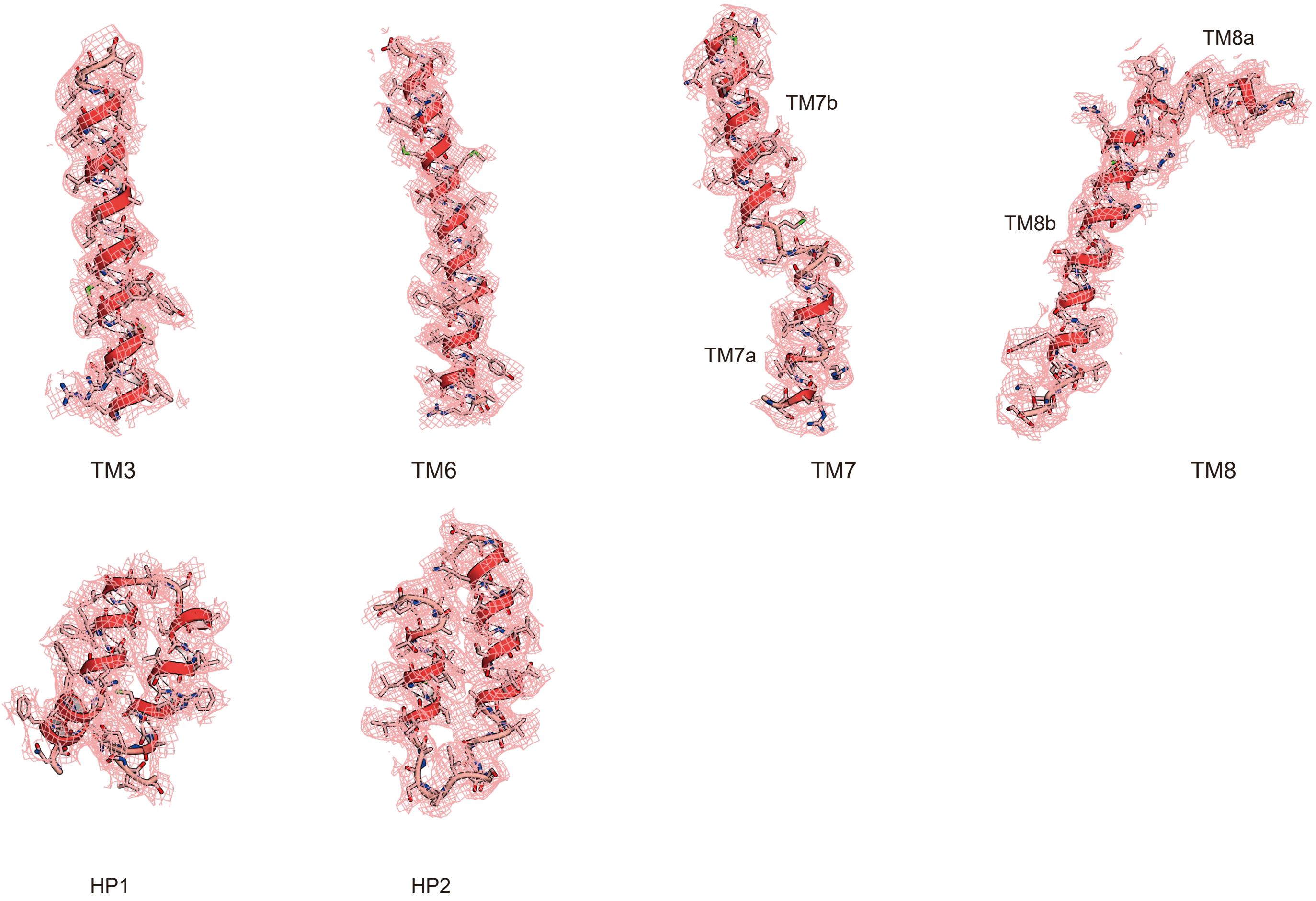


**Supplementary Figure 4 | Atomic model of HsEAAT2 in the cryo-EM map**

(**a**, **b**) Densities of all transmembrane helices and helical hairpins (countoured at 1.0σ). **a**, The densities of the scaffold domain, and **b**, the densities of the transport domain and helical hairpins (HP1 and HP2). TM2, TM5, TM7 and TM8 are each diveded into two segments.

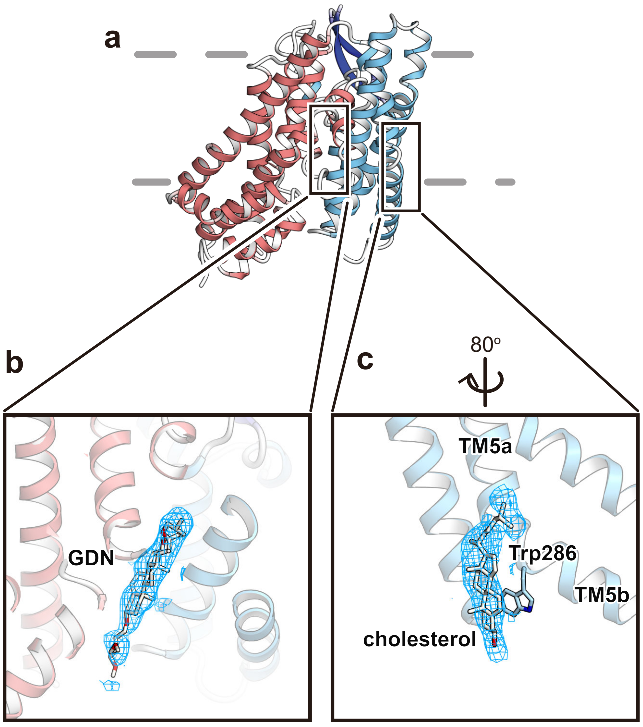


**Supplementary Figure 5│Lipid densities of the HsEAAT2 protomer**

**a**, Overall structure of the HsEAAT2 protomer, viewed from the same orientation as in Fig. 2**a**. **b**, **c**, Close-up views of lipid densities. **b**,Density located between the transport domain and the scaffold domain. **c**, Density located at the scaffold domain. Each density is shown as a light blue mesh.

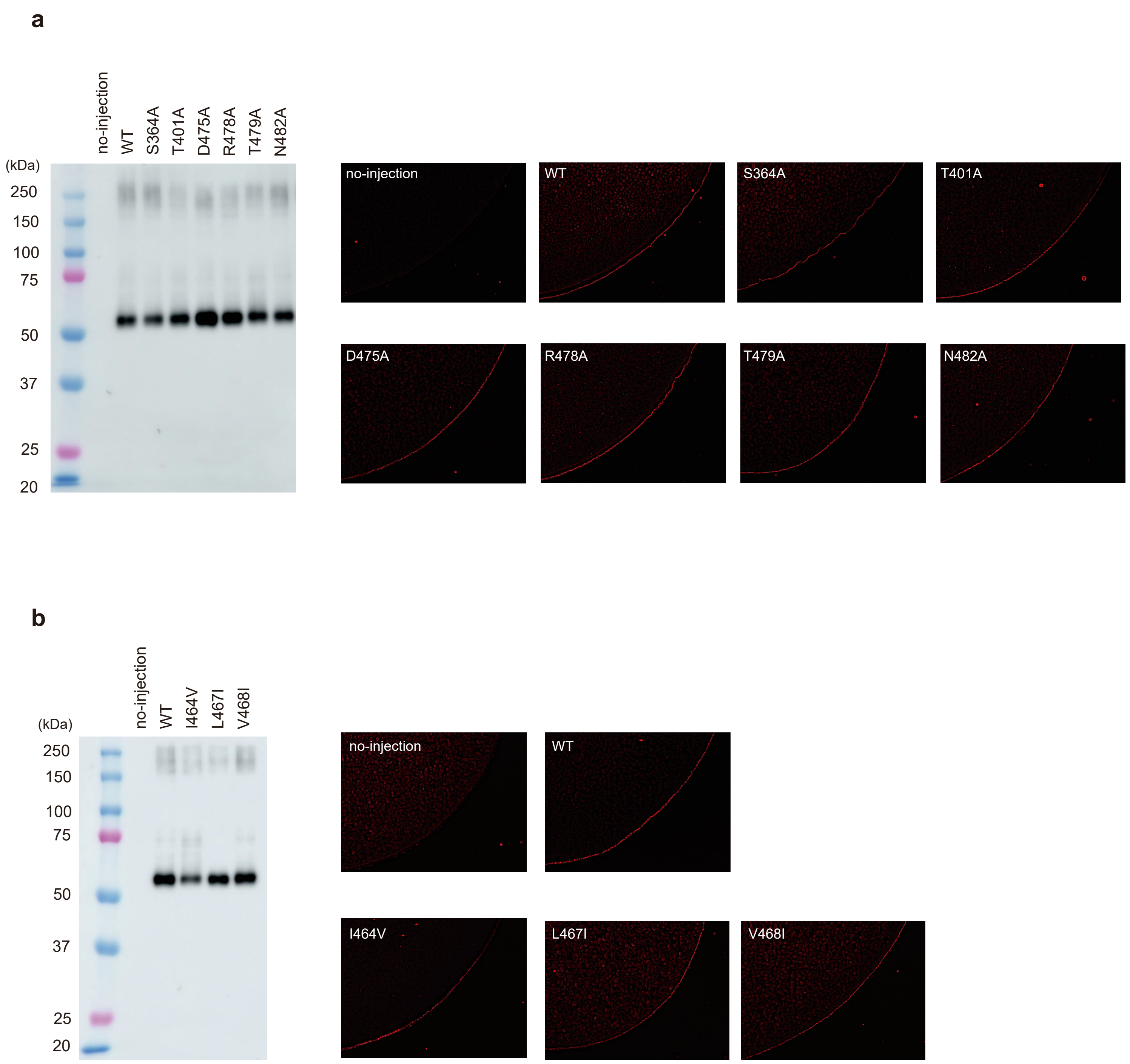


**Supplementary Figure 6 | Expression level and localization of each mutant**

Expression level and localization for **a**) each mutant of the substrate-binding site and **b**) each mutant of the water-accessible cavity. The western blotting analysis and the fluorescence detection at the plasma membrane are shown on the left and right sides, respectively. All measurements were repeated at least three times, and one representative result for each mutant is shown.

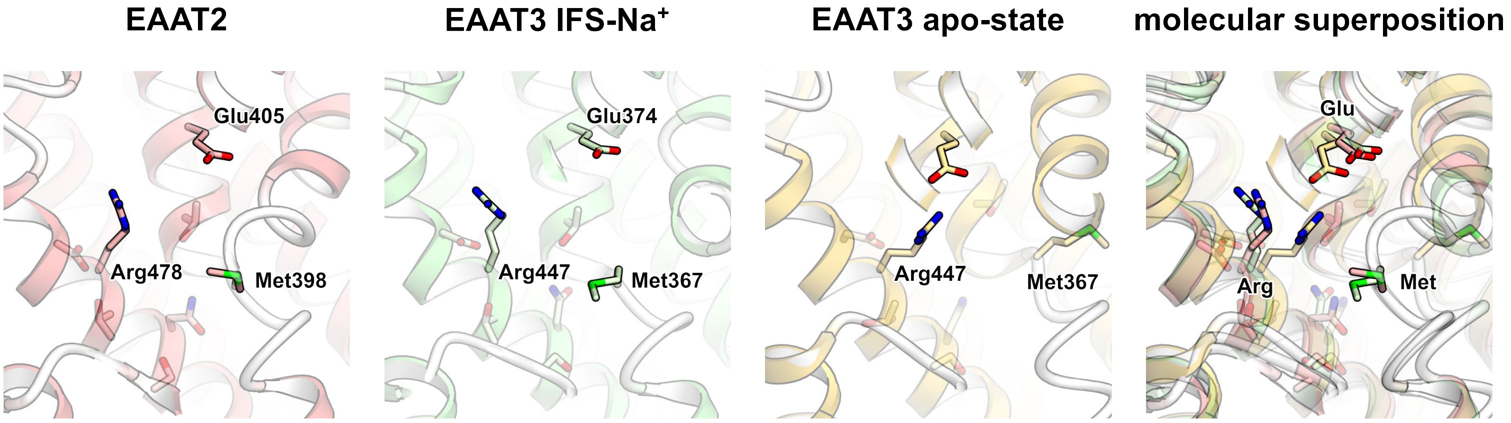


**Supplementary Figure 7 │ Comparison between EAAT2 and EAAT3**

Close-up views of the pockets of EAAT2 substrate-free state, EAAT3 apo-state (PDB ID 6X3F) and EAAT3 IFS-$Na^+$ (6X2Z), and their molecular superposition. Opaque sidechains indicate residues recognizing aspartate and glutamate at the binding site.

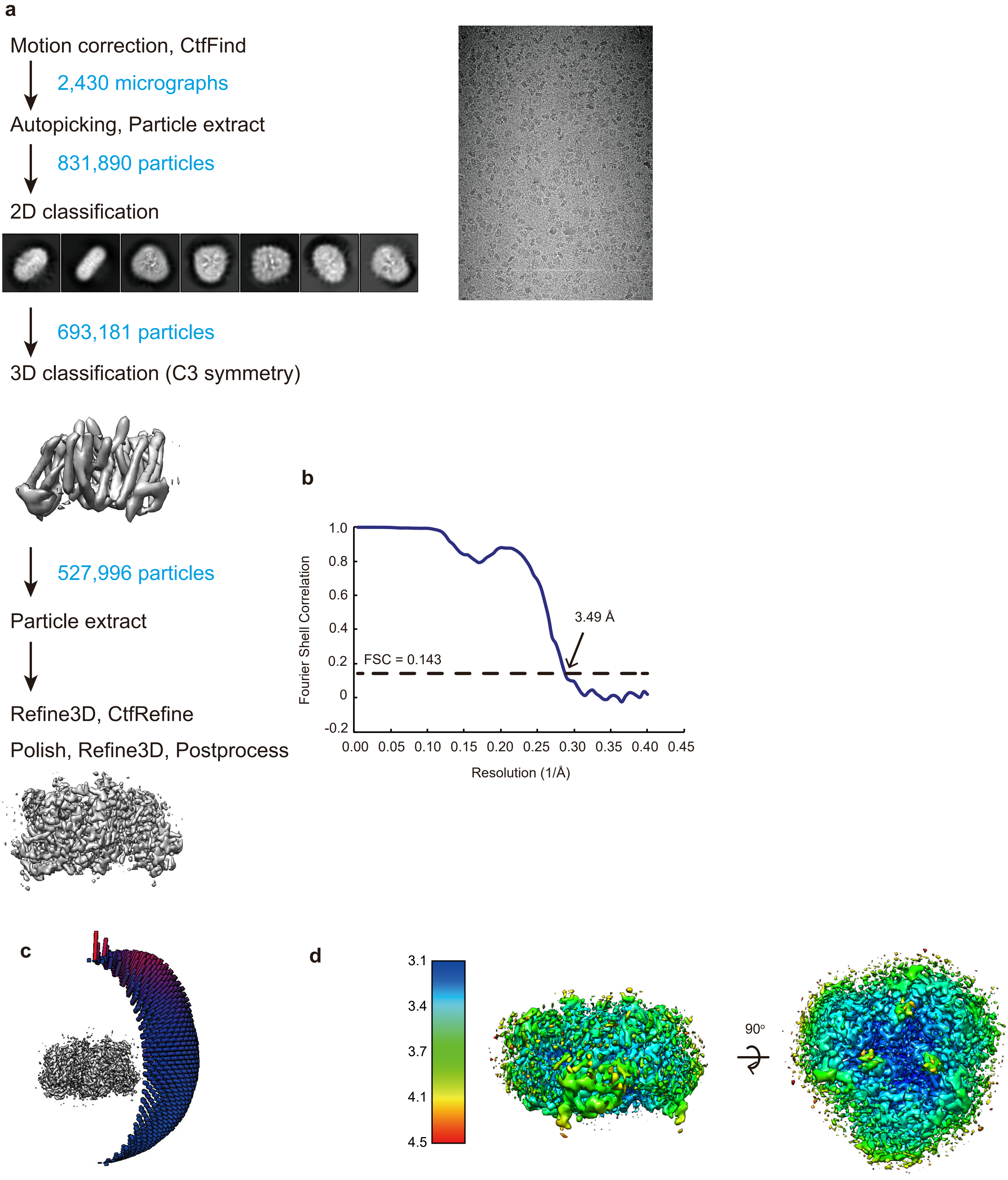


**Supplementary Figure 8│cryo-EM analysis of the HsEAAT2 IFS-WAY213613 state**

**a,** Flow chart of cryo-EM data processing of the HsEAAT2 IFS-WAY213613 state. **b**, Fourier Shell Correlation (FSC) curve of the final 3D reconstruction model calculated using "relion_postprocess" with masked marked 3.49 Å resolution, corresponding to the FSC = 0.143 gold standard cut-off criterion. **c**, Angular distribution plot of particles included in the final 3D reconstruction of the HsEAAT2 IFS-WAY213613 state, with C3 symmetry imposed. **d**, Local resolution of the HsEAAT2 IFS-WAY213613 state.

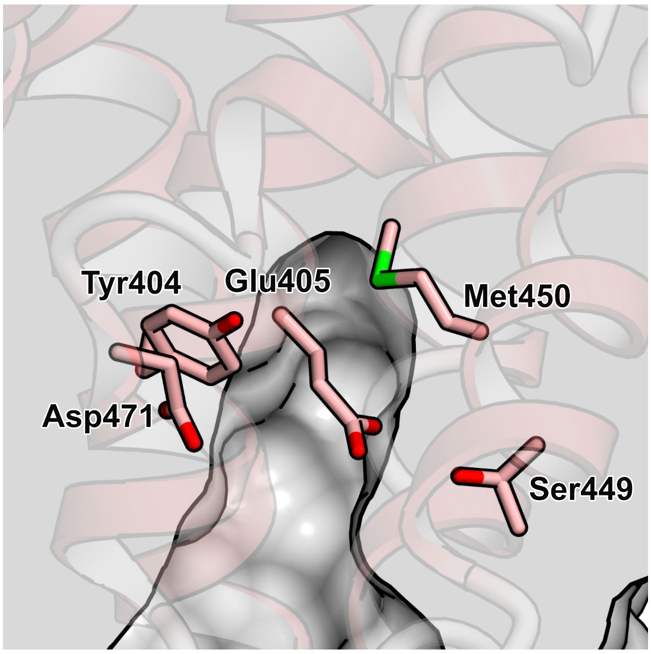


**Supplementary Figure 9│Conserved residues at the cavity**

Close-up view of the cavity formed near the substrate-binding site. Five residues (Tyr404, Glu405, Ser449, Met450 and Asp471) are completely conserved among EAATs.

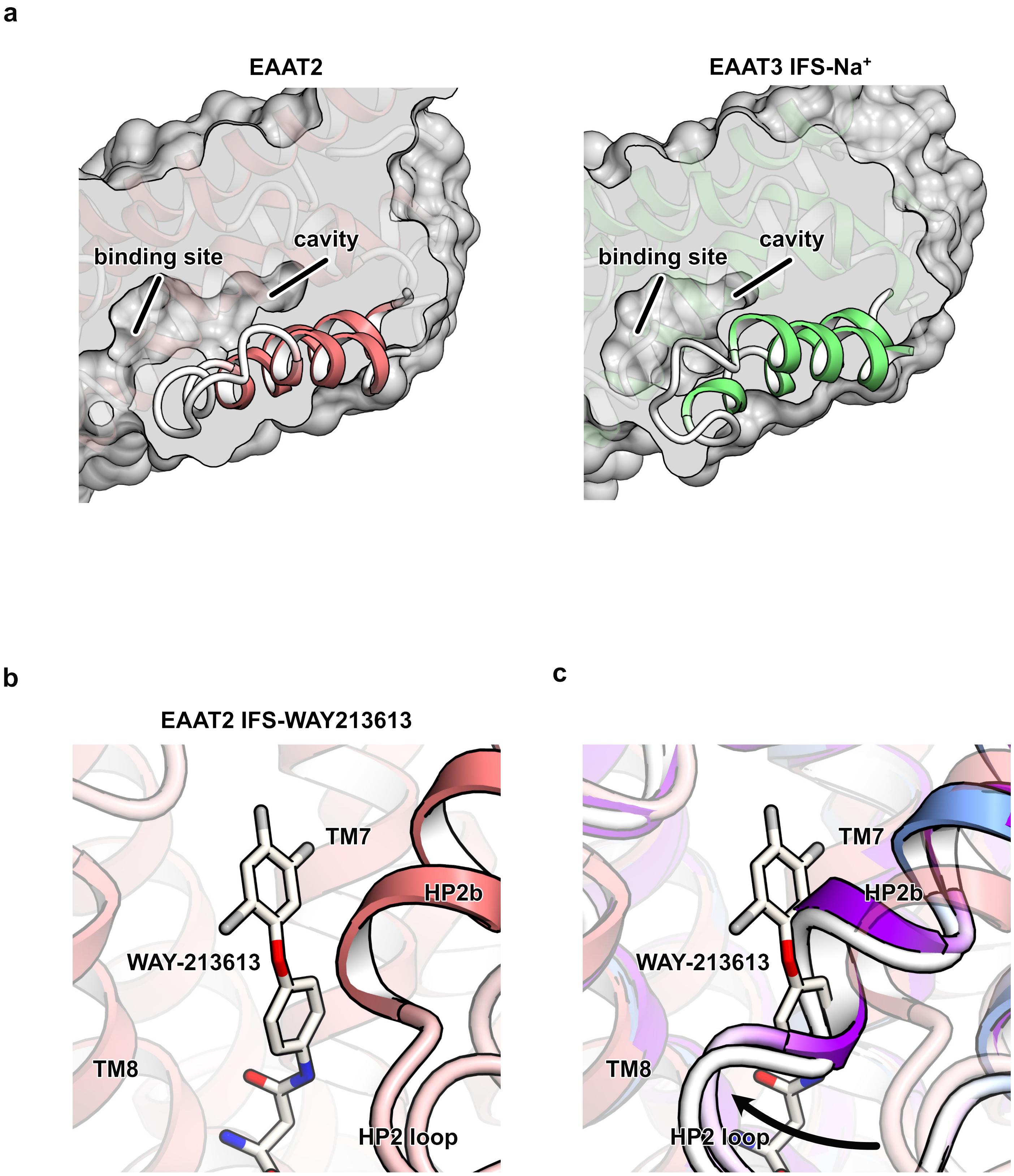


**Supplementary Figure 10 | Structural comparison of transport domains between HsEAAT2 and other SLC1A transporters**

**a**, Cut-away representations of transport domains. The slab thicknesses are the same. **b,** Close-up view of the WAY-213613 binding site. **c**, Molecular superpositions among the IFS-WAY213613 state and Asp-bound states of EAAT1 (purple and PDB ID 5LLU) and EAAT3 (light blue and 6X2Z). The arrow indicates the movement of the HP2 loop.
